# Long-read sequencing of nascent RNA reveals coupling among RNA processing events

**DOI:** 10.1101/230334

**Authors:** Lydia Herzel, Korinna Straube, Karla M. Neugebauer

## Abstract

Pre-mRNA splicing is accomplished by the spliceosome, a megadalton complex that assembles *de novo* on each intron. Because spliceosome assembly and catalysis occur co-transcriptionally, we hypothesized that introns are removed in the order of their transcription in genomes dominated by constitutive splicing. Remarkably little is known about splicing order and the regulatory potential of nascent transcript remodeling by splicing, due to the limitations of existing methods that focus on analysis of mature splicing products (mRNAs) rather than substrates and intermediates. Here, we overcome this obstacle through long-read RNA sequencing of nascent, multi-intron transcripts in the fission yeast *Schizosaccharomyces pombe*. Most multi-intron transcripts were fully spliced, consistent with rapid co-transcriptional splicing. However, an unexpectedly high proportion of transcripts were either fully spliced or fully unspliced, suggesting that splicing of any given intron is dependent on the splicing status of other introns in the transcript. Supporting this, mild inhibition of splicing by a temperature-sensitive mutation in Prp2, the homolog of vertebrate U2AF65, increased the frequency of fully unspliced transcripts. Importantly, fully unspliced transcripts displayed transcriptional read-through at the polyA site and were degraded co-transcriptionally by the nuclear exosome. Finally, we show that cellular mRNA levels were reduced in genes with a high number of unspliced nascent transcripts during caffeine treatment, showing regulatory significance of co-transcriptional splicing. Therefore, overall splicing of individual nascent transcripts, 3’ end formation, and mRNA half-life depend on the splicing status of neighboring introns, suggesting crosstalk among spliceosomes and the polyA cleavage machinery during transcription elongation.

## Introduction

Pre-mRNA splicing results in excision of non-coding introns and ligation of exons to produce mature mRNA. The two-step transesterification reaction involving the 5’ and 3’ splice sites (5’ and 3’ SSs) and the branch point sequence (BPS) is catalyzed by the spliceosome, which must assemble *de novo* on each intron (Wahl et al. 2009). Global short-read sequencing (RNA-seq) of nascent RNA from yeast to human has revealed that most introns are removed from pre-mRNA during transcription by RNA polymerase II (Pol II) (Brugiolo et al. 2013). Therefore, spliceosome assembly and splicing occur on nascent RNA and are closely connected to other processes shaping mRNA expression, from the transcription process itself to 5’ end capping, 3’ end cleavage, and mRNA decay (Bentley 2014; Herzel et al. 2017). For example, 5’ end capping and recruitment of the nuclear cap binding complex promotes spliceosome assembly on the first intron (Gornemann et al. 2005), and histone modifications and variants can influence spliceosome assembly (Gunderson and Johnson 2009; Gunderson et al. 2011; Neves et al. 2017; Nissen et al. 2017). Moreover, splicing and transcription can influence one another to yield splicing-dependent transcriptional pausing and determine alternative splicing patterns (Schor et al. 2013; Dujardin et al. 2014; Fong et al. 2014; Saldi et al. 2016). The mechanisms underlying coordination between RNA processing and transcription are the subject of intense investigation.

Recent efforts have focused on determining the *in vivo* dynamics of gene expression in multiple species and biological contexts, providing insight into the coordination between splicing and transcription. For example, unspliced introns can lead to nuclear retention of 3’ end cleaved transcripts, suggesting that splicing rates can be regulated to promote particular gene expression programs (Bhatt et al. 2012; Boutz et al. 2015; Wong et al. 2016). Over the past ten years, numerous studies have employed a variety of methods – including live cell imaging, metabolic labeling and analysis of nascent RNA – to determine *in vivo* splicing rates; the data indicate a wide range of splicing kinetics that vary according to method and species (Alpert et al. 2017). The core spliceosome itself is highly conserved (Fabrizio et al. 2009); yet, gene architecture is strongly species-dependent and related to splicing mechanisms and likely a key factor defining splicing kinetics. Specifically, the length of introns, their SS and BPS diversity, and intron number per gene increase with the complexity of the organism. Typical human genes contain eight introns (Sakharkar et al. 2005), increasing cellular demands for splicing machinery and generating vast potential for alternative splicing (Lee and Rio 2015). In higher metazoans, introns are ten times longer than exons and typical internal exons are 150 nt long (Zhang 1998), and spliceosomes are thought to assemble following an exon definition mechanism, whereby protein interactions across internal exons are required (Robberson et al. 1990).

When introns are shorter than exons, intron definition triggers spliceosome assembly through bridging of 5’ and 3’ SSs across the intron by the U1 and U2 small nuclear ribonucleoproteins (snRNPs), respectively. The observed high frequency of co-transcriptional splicing in *Saccharomyces cerevisiae* suggests efficient spliceosome assembly through the intron definition mechanism (Alexander et al. 2010; Carrillo Oesterreich et al. 2010; Barrass et al. 2015; Wallace and Beggs 2017). Recently, our lab used two single molecule RNA sequencing methods to analyze the position of Pol II when exon ligation occurs (Carrillo Oesterreich et al. 2016). We found that 50% of splicing events are complete when Pol II is only 45 nucleotides (nt) downstream of 3’SSs, indicating that the spliceosome is physically close to Pol II during catalysis. This raises the important question of how splicing is achieved in an organism that, unlike budding yeast, has multi-intron genes. If splicing occurs as rapidly in multi-intron genes as in single intron budding yeast genes, introns would be spliced in the order of their transcription. However, this has not yet been determined due to a lack of methods that detect the order of intron removal.

Here, we address co-transcriptional splicing efficiency and the possibility that rapid splicing imposes order on intron removal – from first to last intron as transcription proceeds – in *Schizosaccharomyces pombe*, where >1200 protein-coding genes have multiple introns. The *S. pombe* splicing machinery is more similar to higher eukaryotes than to *S. cerevisiae* (Kaufer and Potashkin 2000), and alternative splicing occurs particularly in meiosis (Averbeck et al. 2005; Awan et al. 2013; Bitton et al. 2014; Bitton et al. 2015; Kilchert et al. 2015; Kuang et al. 2017). Splicing rates in *S. pombe* are similar to those reported in *S. cerevisiae* (Alexander et al. 2010; Barrass et al. 2015; Carrillo Oesterreich et al. 2016; Eser et al. 2016). *S. pombe* introns are very short with a median length of 56 nt, whereas exon lengths are similar to higher eukaryotes with an internal exon median of 137 nt (Kupfer et al. 2004; Herzel 2015). This gene architecture is consistent with an intron definition mechanism in *S. pombe* (Romfo et al. 2000; Shao et al. 2012; Fair and Pleiss 2017), suggesting that splicing of individual introns in multi-intron transcripts may occur independently of one other. To globally investigate the progression of splicing during transcription, nascent RNA (nRNA) prepared from proliferating *S. pombe* cells was analyzed by both short- and long-read sequencing (LRS). We report an unexpected coordination between splicing of neighboring introns and polyA site cleavage.

## Results

**Proteomic and transcriptomic characterization of *S. pombe* chromatin**

To isolate nascent RNA, *S. pombe* chromatin was purified and analyzed for RNA and protein composition (Fig. 1A). *S. pombe* cells were harvested in early exponential growth and fractionated into cytoplasm and nuclei (Fig. S1A). Lysed nuclei were further separated into an insoluble chromatin pellet and a soluble nucleoplasmic fraction. Western blot analysis and electrophoresis of nucleic acids was used to evaluate the fractionation (Fig. 1B), showing enrichment of chromatin-associated proteins and genomic DNA and depletion of rRNA in the chromatin fraction. Mass spectrometry confirmed that nuclear, chromatin-associated protein complexes were present, whereas the cytoplasmic fraction contained mainly known cytoplasmic proteins (Table 1, S1B-C, Table S3). Interestingly, many splicing factors were detected in the chromatin fraction, likely reflecting the greater prevalence of intron-containing genes and their cotranscriptional splicing compared to *S. cerevisiae* (Table 1), where splicing factors were not detected on chromatin (Carrillo Oesterreich et al. 2010). Overall, the obtained RNA, DNA and protein signature upon *S. pombe* cell fractionation reflects purified chromatin.

**Figure 1:**
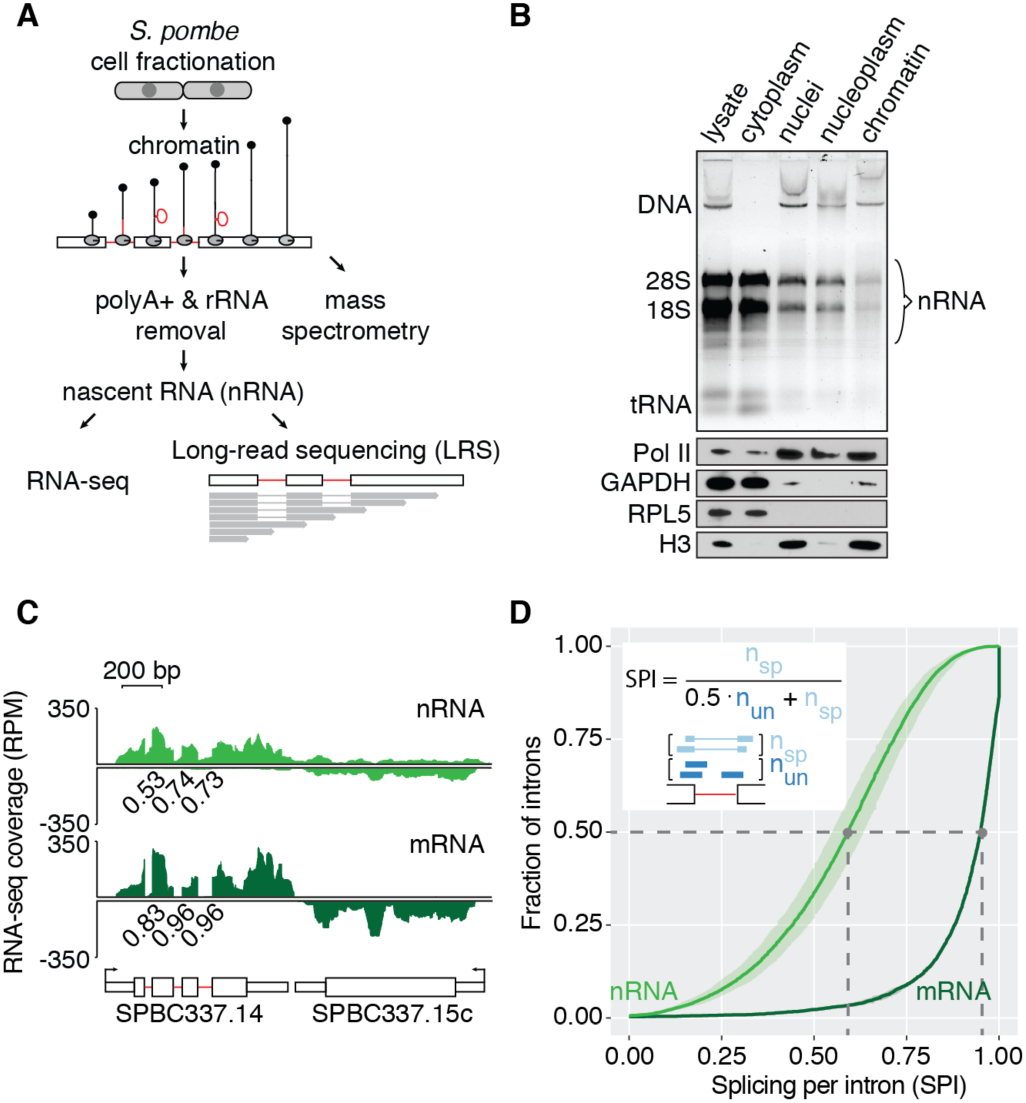
Transcriptome analysis of *S. pombe* chromatin reveals co-transcriptional splicing activity. (A) Schematic of nascent RNA (nRNA) purification from chromatin for short-(RNA-seq) and long-read RNA sequencing (LRS). (B) Enrichment of genomic DNA (DNA) and nRNA in the chromatin fraction and depletion of rRNA (18S & 28S) and tRNA revealed by gel electrophoresis and staining with Gelstar (Lonza). Western blot analysis with antibodies specific for chromatin-associated proteins Pol II and Histone 3 (H3) and cytoplasmic marker proteins GAPDH and RPL5. (C) Nascent and mRNA-seq read coverage (RPM) over a 3-intron gene and a convergent intronless gene. The pooled coverage from 3 biological replicates for each cellular fraction is shown. To assess splicing levels in nRNA and mRNA, splicing per intron (SPI) was calculated from intron junction reads (Herzel and Neugebauer 2015). SPI values are shown for these representative introns underneath the RNA-seq coverage track. (D) Cumulative SPI distribution for *S. pombe* introns (nRNA, n=4,481 introns; mRNA, n=2,181). Mean values (line) with standard deviation (shading) are shown for 3 biological replicates. Grey dashed lines indicate the median splicing levels in the two populations (nRNA 0.59, mRNA 0.95). The inset shows schematizes how the SPI is calculated from the number of ‘spliced’ and ‘unspliced’ junction reads spanning a particular intron.

**Table 1:**
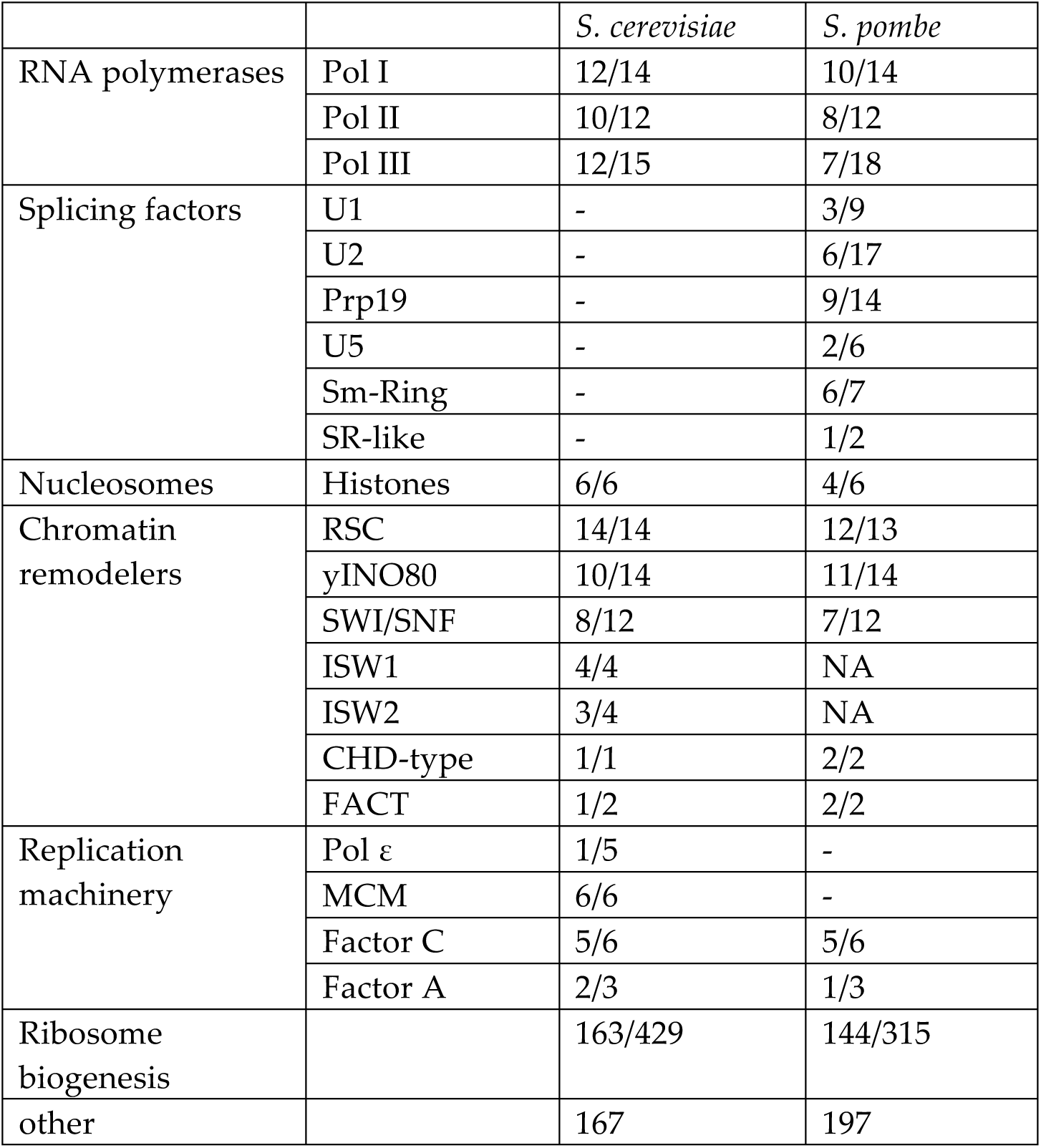
Mass spectrometry of *S. pombe* chromatin with a comparison to *S. cerevisiae*. *S. cerevisiae* data from (Carrillo Oesterreich et al. 2010). 437 proteins were enriched in the chromatin fraction. The table lists the number of detected proteins relative to the number of annotated proteins for a certain function or complex (complete list in Table S3). A dash indicates complete absence of subunits of a protein complex. ISW1/2 are not present in the *S. pombe* genome (NA).

|  |  | *S. cerevisiae* | *S. pombe* |
| --- | --- | --- | --- |
| RNA polymerases | Pol I | 12/14 | 10/14 |
|  | Pol II | 10/12 | 8/12 |
|  | Pol III | 12/15 | 7/18 |
| Splicing factors | U1 | - | 3/9 |
|  | U2 | - | 6/17 |
|  | Prp19 | - | 9/14 |
|  | U5 | - | 2/6 |
|  | Sm-Ring | - | 6/7 |
|  | SR-like | - | 1/2 |
| Nucleosomes | Histones | 6/6 | 4/6 |
| Chromatin remodelers | RSC | 14/14 | 12/13 |
|  | yINO80 | 10/14 | 11/14 |
|  | SWI/SNF | 8/12 | 7/12 |
|  | ISW1 | 4/4 | NA |
|  | ISW2 | 3/4 | NA |
|  | CHD-type | 1/1 | 2/2 |
|  | FACT | 1/2 | 2/2 |
| Replication machinery | Pol $\epsilon$ | 1/5 | - |
|  | MCM | 6/6 | - |
|  | Factor C | 5/6 | 5/6 |
|  | Factor A | 2/3 | 1/3 |
| Ribosome biogenesis |  | 163/429 | 144/315 |
| other |  | 167 | 197 |

To enrich for nascent protein-coding transcripts, RNA purified from the chromatin fraction was further depleted of polyadenylated RNA and rRNA, using bead-based negative selection with oligo-dT and the RiboZero kit (Fig. 1A, S1). Three biological replicates of chromatin-associated, non-polyadenylated RNA and mRNA from the respective cytoplasmic fraction were prepared for RNA sequencing on the Illumina platform, with an average of 14 Mio mapped reads per replicate. As expected, introns and regions downstream of genes showed higher signal coverage in the nascent RNA than in mRNA, indicating the presence of unspliced pre-mRNA and uncleaved, non-terminated RNA (Fig. 1C).

To determine the fraction of spliced introns in nascent RNA and mRNA, we quantified splicing per intron (SPI) values using the fraction of spliced junction reads compared to all junction reads of a particular intron (Herzel and Neugebauer 2015), for the majority of introns (4,481) in the *S. pombe* genome (Figs. 1C-D and S2A; Table S6). An SPI of 1 reflects 100% splicing and an SPI of 0 no splicing of the respective intron. The SPI distribution median for nRNA was 0.59 and for mRNA 0.95. Hence, 50% of introns were spliced to ≥59% in our nRNA dataset. To validate our splicing quantification, 41 introns were randomly selected for semi-quantitative RT-PCR validation, out of which 33 could be evaluated using primers located in the exons flanking the intron of interest (Supplemental methods and Table S2). A strong positive correlation to our nascent RNA-seq data was observed (R=0.7, Fig. S2B), despite the smaller dynamic range of the quantification by RT-PCR. We conclude that *S. pombe* nRNA-seq accurately quantifies co-transcriptional splicing levels for individual introns in the *S. pombe* genome.

### A multitude of intron-specific features correlate with nascent RNA splicing levels

If pre-mRNA splicing of individual introns occurs in the direction of transcription, a decrease of global pre-mRNA splicing levels towards 3’ ends of genes might be expected. Global nRNA splicing levels did not change in the direction of transcription, but rather with the relative distance to the transcript start and end, generally showing higher splicing for internal introns (Fig. 2A). The group of single, first and last introns were spliced to a lesser extent than internal introns, which showed similar SPIs independent of their position (median SPI 0.62). This trend was also detected in cytoplasmic mRNA (Fig. 2B), suggesting that incompletely spliced transcripts can be exported to the cytoplasm. To test if terminal introns show generally lower splicing in individual genes, the difference of SPIs of adjacent introns was calculated (SPI of 5’ intron – SPI of 3’ intron). 25% of those intron pairs showed significant differences in splicing levels (Fig. 2C). Consistent with the earlier observation of lower first intron splicing, 176 first introns were spliced less (52% of 5’ less group) and only 73 were spliced more (27% of 5’ more group) than the second intron. Next, we compared intron features between the more and less frequently spliced introns in the pairs with significant splicing differences. The 42 analyzed intron features included RNA-seq-derived measures, e.g. SPIs and coverage of exons and introns in RPKM, and gene architecture measures of exons and introns, e.g. splice site and BPS strength, length, conservation in the fission yeast clade and GC-content (see Supplemental methods and Table S6, S9 for a complete overview of intron features). Fifteen features, e.g. cytoplasmic mRNA-seq intron coverage and 5’SS and BPS strength, were significantly associated with low or high splicing in an intron pair (Fig. 2D). Interestingly, mRNA expression also correlated with nascent RNA splicing, with the highest spliced introns belonging to mRNAs with the highest levels (Fig. S2C). We conclude that nRNA intron splicing does not decrease in the direction of transcription, but first, last and single introns tend to be less co-transcriptionally spliced than internal introns.

**Figure 2:**
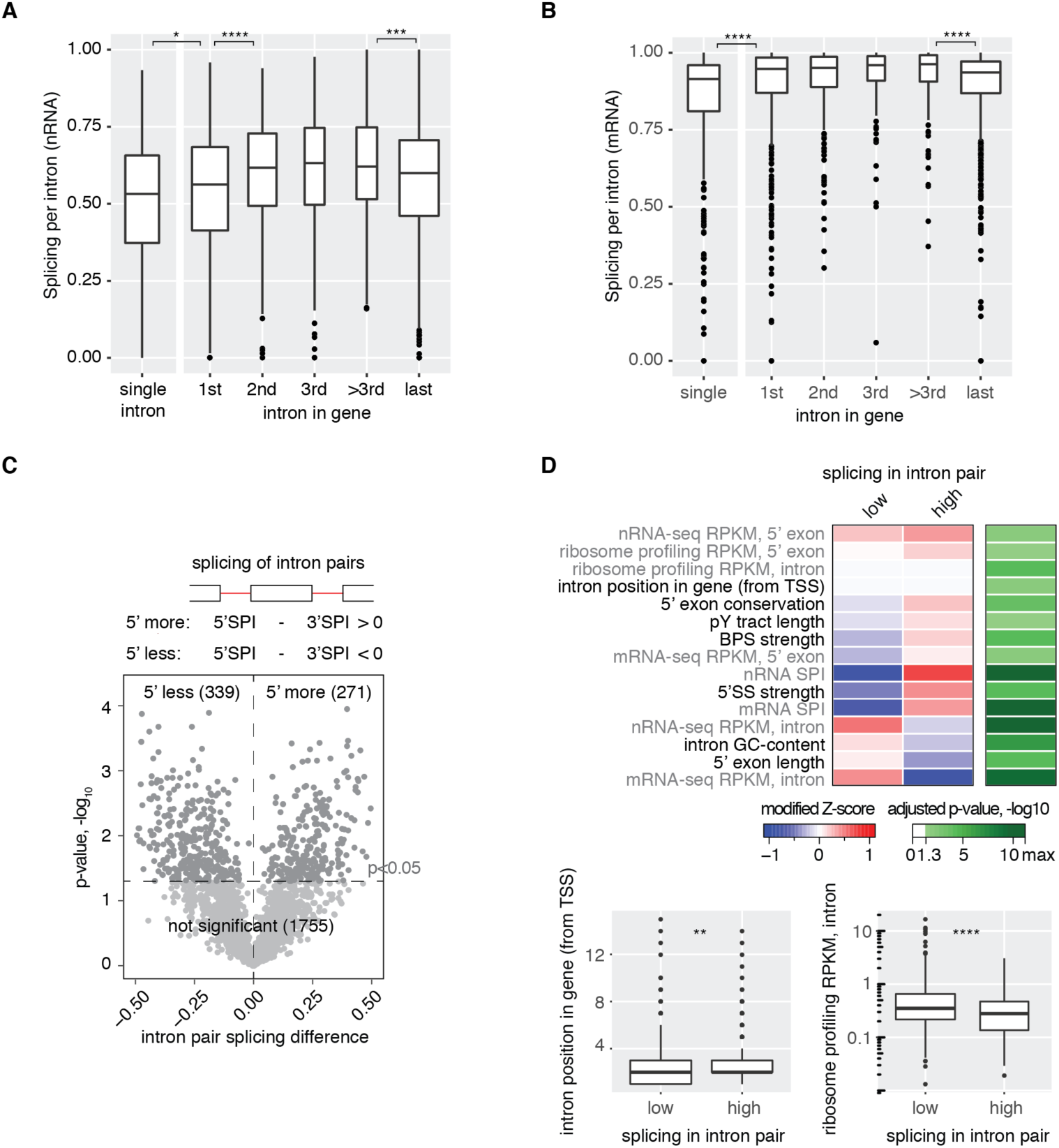
Pre-mRNA features that correlate with the extent of co-transcriptional intron splicing. (A) Co-transcriptional splicing levels differ between introns in different gene positions. The boxplot shows the distribution of nascent RNA SPIs for the group of single intron genes and first, internal (2^nd^, 3^rd^ and other) or last introns in multi-intron genes. The boxwidth corresponds to the respective group size. (B) mRNA splicing levels differ between introns in different gene positions. The boxplot shows the distribution of mRNA SPIs for the group of single intron genes and first, internal (2^nd^, 3^rd^ and other) or last introns in multi-intron genes. The boxwidth corresponds to the respective group size. (C) One quarter of introns are significantly less or more spliced than the next downstream (3’) intron in nRNA. This is depicted as volcano plot, summarizing data from 3 biological replicates. (D) 15/42 analyzed gene architecture features correlate significantly with differentially spliced intron pairs (sequence-based in black font and RNA-seq derived in grey font). The smaller intron position for “low” spliced introns in a pair (first intron – 1, second intron – 2 etc.) is consistent with enrichment of first introns in the “5’ less spliced” group. The median modified Z-score is shown for each feature with significant difference between the “low” and “high” groups and the respective negative log10 of the Bonferroni-corrected p-value is given. For two features, no change in the median modified Z-score is visible. The respective feature distribution is thus presented as boxplot below. Asterisks indicate significance of direct neighbors according the Wilcoxon-rank sum test (p < 0.05 *, p < 0.01 **, p < 0.001 ***, p < 0.0001 **** after Bonferroni-correction) in A, B & D.

### Full-length nascent RNA sequencing of multi-intron transcripts

To assess the order of intron removal directly, nascent RNA was converted into double-stranded cDNA for long-read sequencing (LRS) on the Pacific Biosciences platform. A DNA adaptor with five random nucleotides at the 5’ end was ligated to the 3’ end of all nascent RNAs. This diminishes possible ligation biases from specific 3’ nucleotides and preserves single molecule information (Zhuang et al. 2012; Herzel 2015; Mayer et al. 2015). Template-switching reverse transcription enabled the generation of full-length double-stranded cDNA by PCR, by attaching a universal sequence to the 5’ end. The inclusion of sample-specific barcodes allowed pooling of cDNA preparations, prior to Pacific Biosciences library preparation (Fig. S4). A total of 8 SMRT cells yielded 169,000 high-quality, non-polyadenylated, full-length transcripts from two biological replicates (Supplemental methods, Fig. S4C-G, Table S4). 21% of those transcripts overlapped with intron-containing genes with a median of 7 transcripts per gene. Transcript counts per gene correlated well with expression data from nascent RNA-seq (Fig. S4F).

Full-length transcript sequence informs on the history of transcription and splicing of every RNA molecule. Nine examples from multi-intron genes and full-length nascent transcripts detected by long-read sequencing are shown in Fig. 3A. Most transcript 5’ ends mapped close to the annotated TSS or to well-defined positions downstream of the existing annotation (e.g. SPAC16E8.15). 3’ ends correspond to the nucleotide position of elongating Pol II at the time of cell lysis. As seen previously for single intron nascent RNAs, introns were often spliced out when Pol II was positioned just downstream of the intron (Carrillo Oesterreich et al. 2016) (Fig. 3A, S4H). To evaluate the order of intron removal, it was necessary to identify partially spliced transcripts. Transcripts overlapping multiple introns were classified as “all spliced” if all introns were removed, as “all unspliced” if all introns were present, or as “partially spliced” if some but not all of the introns were spliced (see Fig. 3A). Partially spliced transcripts in the same gene were often associated with a specific intron being (un)spliced, e.g. unspliced intron 1 in SPBC428.01c or the unspliced intron 1 in SPBP8B7.11.

**Figure 3:**
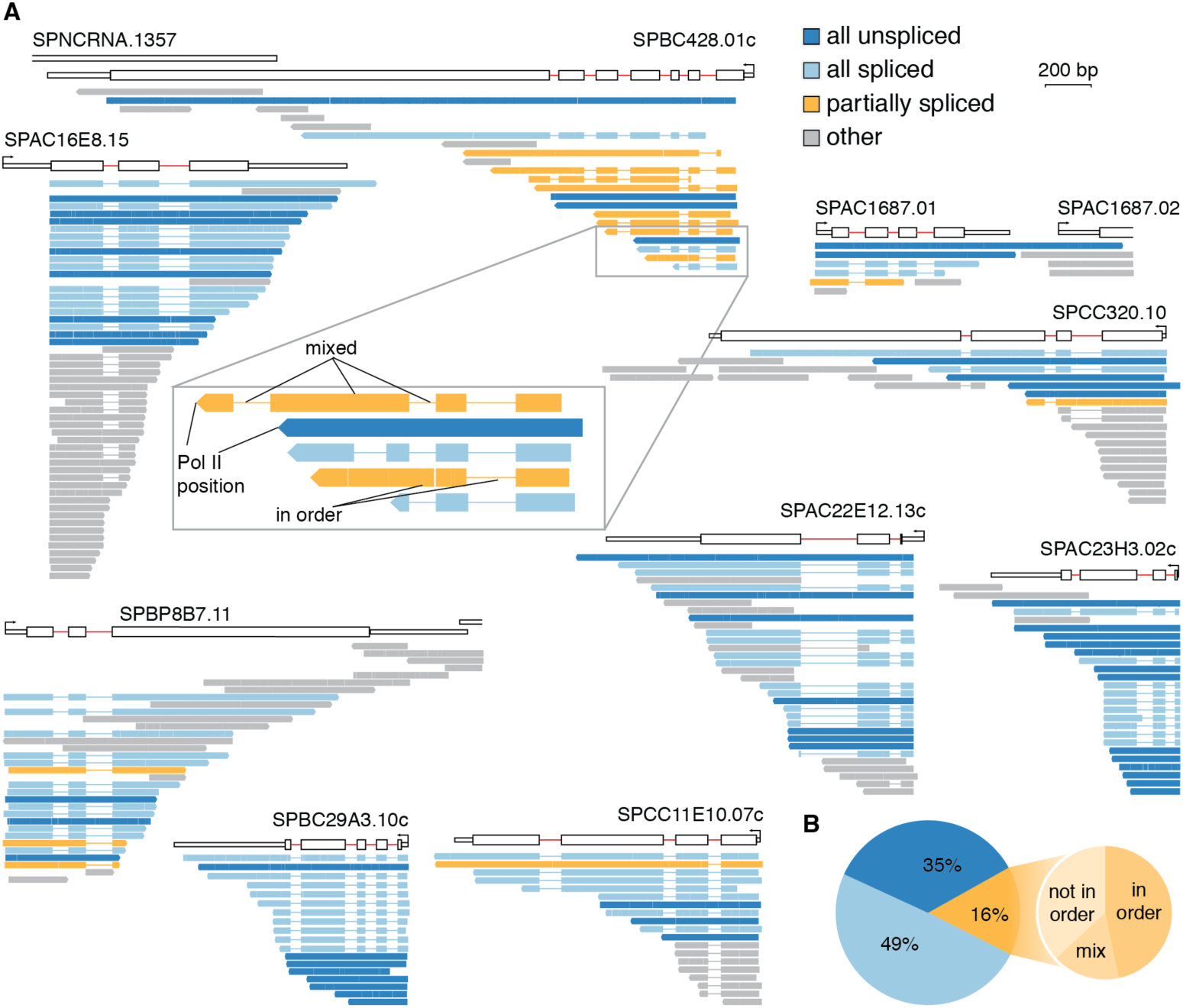
Single molecule long-read sequencing reveals predominantly “all or none” splicing of multi-intron transcripts. (A) Full-length transcripts mapping to nine genes with more than one intron are shown underneath each gene diagram. Transcripts are color-coded according to their splicing profile: “all unspliced” (dark blue), “all spliced” (blue) and “partially spliced” (orange). Thin lines in transcripts indicate that the intron sequence is absent (due to splicing). Transcripts overlapping with <2 introns (“others”, grey) cannot be used to analyze the order of intron splicing. The inset shows five transcripts of gene SPBC428.01c, highlighting two subclasses of partially spliced transcripts and that transcript 3’ ends refer to Pol II positions during transcription. (B) All unspliced, all spliced and partially spliced fraction in the transcriptome. The large pie chart shows the proportion of the three splicing categories from nRNA LRS. The small pie chart depicts the fraction of partially spliced transcripts, which show intron removal ‘in order’ (all introns are spliced upstream an unspliced intron in a particular transcript, compare zoom in of SPBC428.01c in A), ‘not in order’ (at least one intron is unspliced upstream of a spliced intron in a particular transcript) or a mixed pattern (compare inset of SPBC428.01c in A).

A comparison of the partially spliced transcripts identified by LRS to our nRNA-seq data showed a highly significant overlap between datasets (p-value=4*10^-08^, Fisher’s exact test). 5’ introns classified as ‘unspliced’ or ‘spliced’ in partially spliced transcripts distributed according to the difference in nRNA-seq splicing values between adjacent intron pairs and thus recapitulated the differences in SPI by nRNA-seq (Fig. S5A). To analyze partial splicing patterns with respect to the direction of transcription, partially spliced transcripts were classified as ‘in order’ (5’ introns spliced and one or more 3’ introns unspliced), ‘not in order’ (at least one unspliced intron followed by at least one spliced intron) or transcripts with ‘mixed’ pattern that contained one or more unspliced intron(s) upstream of a spliced intron and downstream of a spliced intron (Fig. 3B, middle panel). Thus, ‘mixed’ transcripts combine both ‘in order’ and ‘not in order’ splicing. The fraction of ‘in order’ and ‘not in order’ transcripts was similar, suggesting that ordered intron removal is not strictly enforced. This is consistent with the nascent RNA-seq data, which did not show a splicing decrease from 5' to 3', but lower splicing of first and last introns. Intron feature analysis showed that 5’SS and BPS strength were reduced in unspliced introns compared to spliced introns of partially spliced transcripts, as well as the nRNA SPIs (Fig. S5B). Further, unspliced introns were associated with higher RNA-seq coverage in our cytoplasmic mRNA-seq data set. Comparison of our nascent LRS and published mRNA LRS data showed a decrease of partially spliced transcripts to 43% from nascent RNA to mRNA (Fig. S5C), indicating that a substantial fraction of partially spliced transcripts can be polyadenylated and exported to the cytoplasm, reflecting examples of intron retention in *S. pombe*. In conclusion, LRS of nRNA provides a snapshot of splicing states of multi-intron transcripts. The subcategory of partially spliced transcripts recapitulates the splicing differences seen by nRNA-seq and carries signatures of gene-specific intron retention.

### “All or none” splicing of single transcripts predominates

The number of partially spliced transcripts (697) for the above analysis of splicing order was surprisingly small. This caused us to consider how many partially spliced transcripts would be expected from our dataset. Given that each spliceosome assembles anew on every intron in each transcript and that *S. pombe* introns are mainly defined by intron definition, we expected that pre-mRNA splicing events of neighboring introns would be independent of one another. This is illustrated in a prediction for all expressed *S. pombe* intron-containing genes with more than one intron (Fig. 4A), utilizing the intron splicing frequencies from our nascent RNA-seq data. In this scenario, 81% of transcripts would be either partially or completely spliced and only 19% completely unspliced. Surprisingly, this prediction was not recapitulated by our LRS data, in which 84% of transcripts were either completely spliced or unspliced. Only 16% of transcripts were partially spliced, 2.8 times less than expected if splicing of adjacent introns were independent of each other (Fig. 4A).

**Figure 4:**
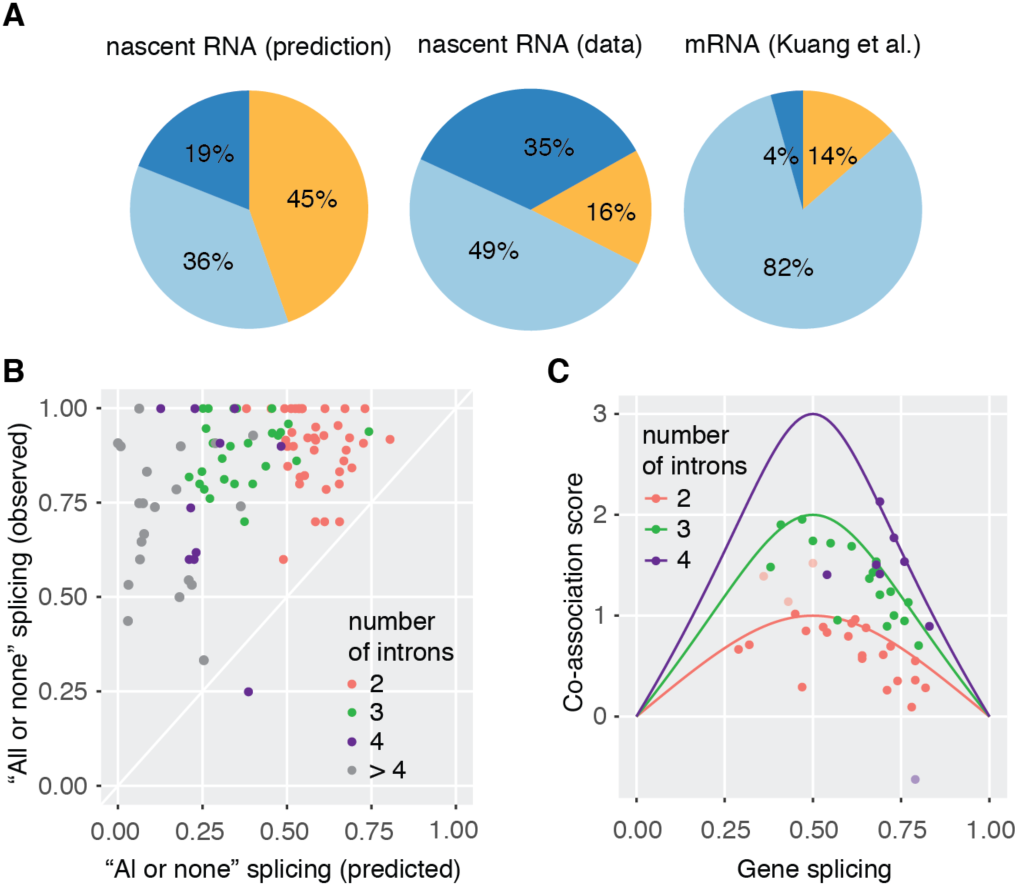
High degree of splicing co-association in multi-intron transcripts. (A) Predicted and observed “all unspliced”, “all spliced” and “partially spliced” fractions in the transcriptome. Left panel, prediction of splicing categories from nRNA-seq, assuming independence of splicing of adjacent introns. Predicted values were calculated for the first two introns of genes with two and more introns for nRNA (see Supplemental methods for details). Middle panel, proportion of the three splicing categories from nRNA LRS as in 3B. Right panel, proportion of the three splicing categories from total mRNA LRS (data from (Kuang et al. 2017)). (B) Predominant “all or none” splicing in multi-intron genes. The predicted fraction of “all or none” splicing (fraction of all spliced or unspliced transcripts calculated from nRNA-seq, assuming splicing independence) is plotted against the observed fraction of “all or none” splicing for 100 genes with 10 or more sequenced transcripts by LRS. (C) Co-association of splicing is close to the maximal possible co-association value. A co-association score was calculated as the log2-fold change of the observed (LRS) to predicted (RNA-seq) fraction of “all or none” splicing. The maximal possible co-association score for 2, 3 and 4-intron genes was calculated (log2(1/gene splicing)) and plotted versus the mean gene splicing value (solid line). Co-association scores for individual genes fall below or on top of this line, suggesting maximal co-association for most genes (median 86%, calculated as ratio of observed co-association score over maximal co-association score at particular gene splicing). Outliers are indicated as lighter points and can be explained by inconsistencies in gene annotation.

To challenge the robustness of this finding, a similar analysis was carried out on a gene-by-gene basis for 100 genes with 10 or more transcripts spanning multiple introns. For 99 of those genes the observed fraction of “all or none” splicing was greater than the predicted fraction (Fig. 4B). To estimate the degree of co-association between individual splicing events, a co-association score was calculated, which we defined as the log2-fold change of observed and predicted fraction of “all or none” transcripts (Fig. 4C, S5E). Note that the maximal co-association score increases with increasing intron number (Fig. S5F). Most genes had co-association scores close to the maximal possible score (Fig. 4C, S5G). Hence, we conclude that partial transcript splicing is the exception in *S. pombe* and suggest that most introns are removed cooperatively. Interestingly, the number of introns in the transcript does not influence the gene-by-gene tendency for co-association (Fig. S5G). Here, the combined analysis of short- and long-read nascent RNA sequencing data enabled us to distinguish between completely spliced mRNAs, mRNAs with retained introns, or completely unspliced RNAs. Our data indicate that splicing in *S. pombe* occurs soon after introns are fully transcribed and may stimulate splicing of further downstream introns, resulting in “all or none” splicing of transcripts with multiple introns.

### Mild global splicing inhibition increases “all unspliced” transcripts

The above findings suggest that the presence of an unspliced intron might negatively impact the splicing of neighboring introns within the same transcript. To test this hypothesis, we profiled nascent RNA splicing in the temperature-sensitive mutant *prp2-1* (homolog of mammalian U2AF65, involved in 3’SS recognition), which is known to reduce mRNA splicing levels upon shifting to the non-permissive temperature (Sridharan et al. 2011; Lipp et al. 2015). By semi-quantitative RT-PCR, no splicing differences between wild-type (WT) and *prp2-1* cells were detected at the permissive temperature (Fig. 5A, S6A). After a two-hour shift to the non-permissive temperature, a ∼50% reduction in pre-mRNA splicing was observed for five analyzed introns in *prp2-1* relative to WT in both cytoplasmic and chromatin fractions (Fig. 5A, S6A-B). LRS was performed on nRNA isolated from *prp2-1* and WT cells in three biological replicates. Similar to the RT-PCR experiment, splicing levels were reduced by half in *prp2-1* (Fig. 5B-C). Global differences in transcription of intron-containing genes were not detected between the two stains (Fig. S6E). Classification of transcripts into “all unspliced”, “all spliced” and “partially spliced” transcripts revealed that the proportion of “all unspliced” transcripts increased from 40 to 59% in *prp2-1* and had only a minor impact on the fraction of “partially spliced” transcripts (Fig. 5B, D). This is consistent with an inhibitory effect of individual unspliced introns on the splicing of the other introns in the same transcript. This is supported by adjusting the prediction from nRNA-seq to match the degree of global splicing inhibition seen in the *prp2-1* LRS data (Fig. S6C). Finally, we asked whether the *prp2-1* mutation affected mRNA levels. If unspliced transcripts were predominantly degraded, the relatively mild inhibition of co-transcriptional splicing by *prp2-1* should specifically reduce mRNA levels for intron-containing genes. Indeed, reanalysis of *prp2-1* and WT mRNA expression data measured by microarrays in (Lipp et al. 2015) showed stronger downregulation of mRNA levels for intron-containing genes than intronless genes at the non-permissive temperature for *prp2-1* (Fig. 5E). We conclude from these data that the *prp2-1* mutation increases the levels of “all unspliced” nascent transcripts, consistent with a pervasive inhibitory effect on the splicing of the other introns in the same transcript, and ultimately leads to downregulation of mRNA levels.

**Figure 5:**
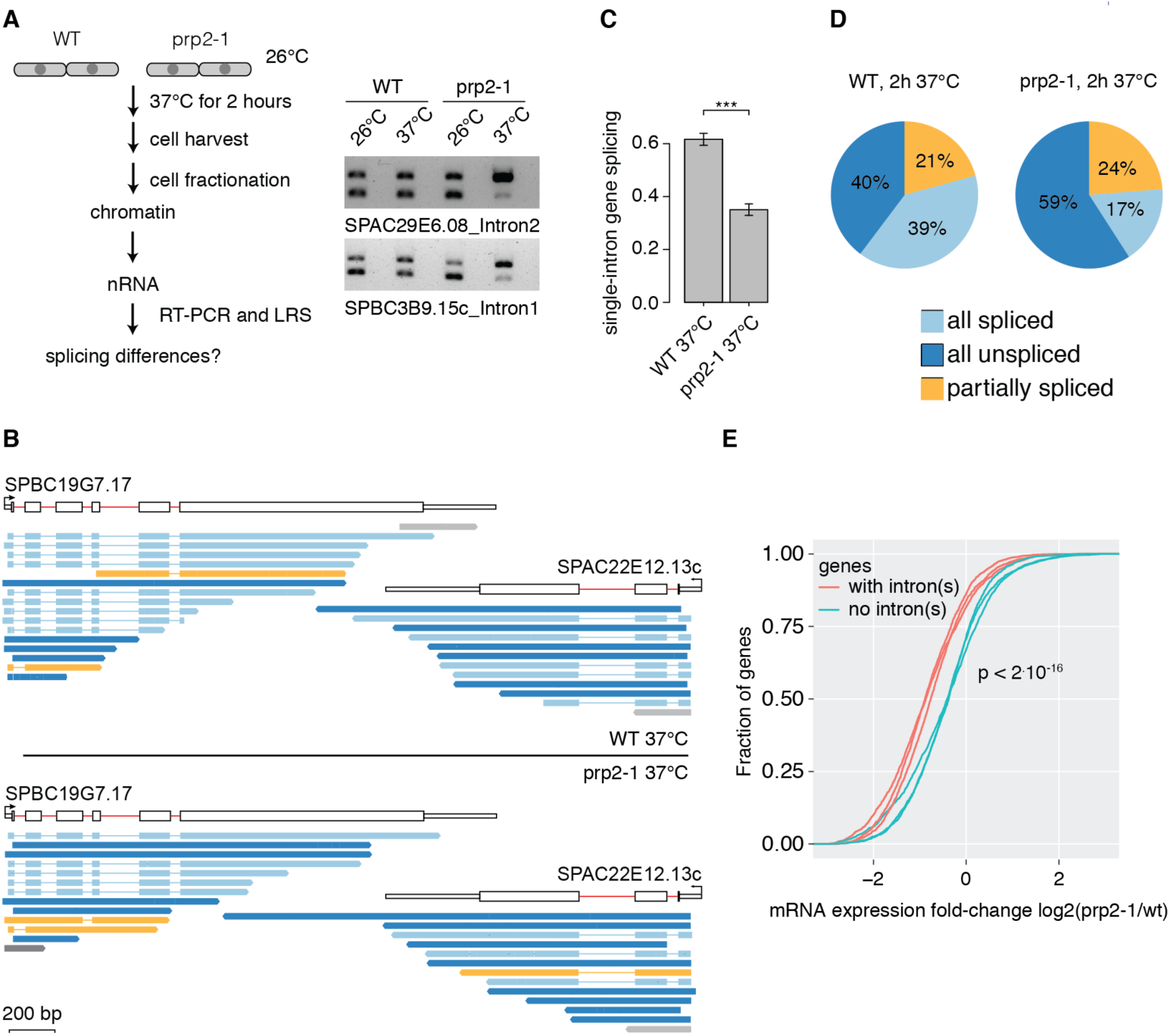
Inhibition of pre-mRNA splicing increases “all unspliced” transcript levels and reduces mRNA levels. (A) Left panel, schematic of the workflow to generate splicing profiles in the temperature-sensitive *prp2-1* mutant and WT strains at non-permissive temperatures. Right panel, RT-PCR shows increased levels of unspliced nascent RNA in the *prp2-1* mutant after 2 hours of growth at 37°C for 2 introns compared to the WT *972h-* strain. –RT control was loaded in lanes (empty) adjacent to the +RT samples. (B) Nascent RNA WT and *prp2-1* LRS read coverage over 2 genes. (C) Barplot indicating splicing frequency for single intron genes in *prp2-1* relative to WT at 37°C calculated from LRS data. The standard deviation from 3 biological replicates is given and asterisks indicate significance according the Student’s t-test (p < 0.001 ***). (D) Comparison of the “all unspliced”, “all spliced” and “partially spliced” fractions in the transcriptome in WT and *prp2-1*. (E) Reduced mRNA levels for intron-containing genes compared to intronless genes in *prp2-1* at 37°C. Cumulative distribution of expression changes between *prp2-1* mutant and WT for intron-containing and intronless genes (data from 3 replicates from (Lipp et al. 2015), p-value from Kolmogorov-Smirnoff test between intronless and intron-containing group).

### Failure to splice co-transcriptionally reduces cytoplasmic mRNA levels

Downregulation of mRNA levels caused by degradation of unspliced pre-mRNAs suggests a possible mechanism to fine tune gene expression through co-transcriptional splicing. To determine whether this is a global phenomenon in cells, genes were grouped into two classes following nRNA-and mRNA-seq expression quantification using cufflinks: 1) genes with mRNA levels lower than nRNA levels and 2) genes with mRNA levels higher than or similar to nRNA levels. Higher mRNA levels than nRNA levels can arise in cases where the mRNA half-life is long (Fig. S6F). Identical mRNA and nRNA levels suggest that transcription levels define mRNA expression. If mRNA levels are lower than nRNA levels, RNA decay of pre-mRNA reduces the fraction of mRNA. For group 1 genes, the average pre-mRNA splicing levels were significantly lower than for group 2 genes, underlining the contribution of unspliced RNAs to establish mRNA expression levels (Fig. 6A).

**Figure 6:**
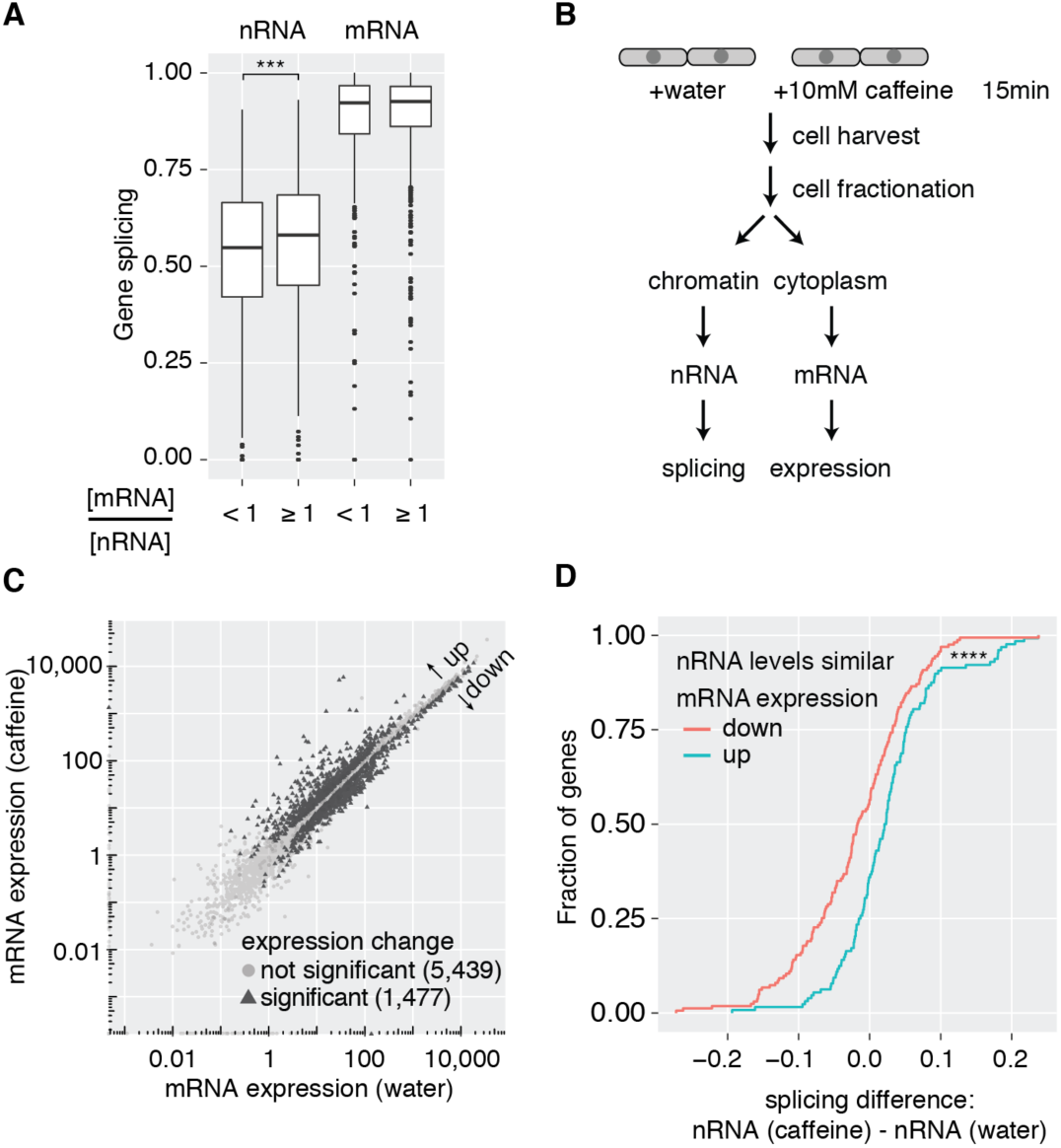
Co-transcriptional splicing correlates with higher mRNA levels. (A) Boxplot of nRNA and mRNA gene splicing levels after grouping according to cytoplasmic mRNA levels relative to nRNA levels (p-value p < 0.001 ***, Wilcoxon-rank sum test). (B) Experimental outline to induce changes in gene expression upon caffeine treatment in *S. pombe* cells. (C) Correlation of mRNA expression values between the two conditions identifies 1,477 differentially expressed genes (FDR-adjusted p-value < 0.05, FDR <= 0.05; 566 of those are intron-containing). (D) Cumulative distribution of nRNA gene splicing differences between caffeine treatment and control. Only genes without significant changes in nRNA levels, but significant differences in mRNA expression were considered (p < 0.0001 ****, Kolmogorov-Smirnoff test between ‘mRNA expression up’ (n=52) and ‘mRNA expression down’ (n=65) gene group).

Next, we sought to determine the contribution of splicing inhibition to physiologically relevant cell signaling and shifting gene expression programs. To alter mRNA levels globally, exponentially growing *S. pombe* cells were treated with 10 mM caffeine for 15 min (Fig. 6B), which elicits a cellular response similar to nitrogen starvation (Rallis and Bahler 2013). Gene expression and splicing profiles were quantified by nRNA-seq and cytoplasmic mRNA-seq for three biological replicates. In agreement with a previous study, ∼1,500 mRNAs changed levels significantly after treatment (Fig. 6C, S7A). No significant global splicing changes were detected when grouping genes according their respective expression profiles in nRNA and mRNA (Fig. S7B). However, that analysis did not account for potential changes in transcription. To determine if co-transcriptional splicing levels alone correlate with mRNA levels, we identified genes among the up- and down-regulated mRNAs whose nascent RNA levels did not change with treatment. Interestingly, significantly lower co-transcriptional splicing frequencies were observed for the down-regulated mRNAs (Fig. 6D), indicating that failure to splice co-transcriptionally leads to lower gene expression.

### Coordination of splicing status, polyA site cleavage, and nuclear degradation

Strong links between splicing and polyA site cleavage (Herzel et al. 2017) led us to determine whether the persistence of unspliced introns in nascent RNA might impact downstream RNA processing events such as polyA site cleavage. Indeed, 574 genes in our WT datasets contained unspliced introns and had 3’ ends located downstream of annotated polyA cleavage sites (Fig. 7A-C). The majority of intron-containing transcripts that had a 3’ end after the annotated polyA cleavage site were completely unspliced (Fig. 7B). Nine out of ten of these instances were validated by RT-PCR (Fig. S8A). Conversely, 95% of polyadenylated transcripts were spliced (Fig. S8D). Given our previous evidence that unspliced transcripts are preferentially degraded (see above), we considered it likely that unspliced transcripts extending over the polyA site might be degraded rather than further processed. Indeed, total RNA-seq data from a nuclear exosome subunit deletion strain (*Δrrp6*) showed a general increase in the amount of unspliced RNA and higher sequencing read coverage downstream of polyA sites (Fig. 7D, Fig. S8B) (Zhou et al. 2015). For the high coverage gene SPAC1805.11c, the signal downstream of the polyA site was substantially higher in the *Δrrp6* strain than in WT, measured by RT-qPCR (Fig. 7E). This finding substantiates the notion that splicing cooperativity among introns extends to coordination with polyA site cleavage, highlighting a pathway to mRNA degradation based on reduced co-transcriptional splicing efficiency.

**Figure 7:**
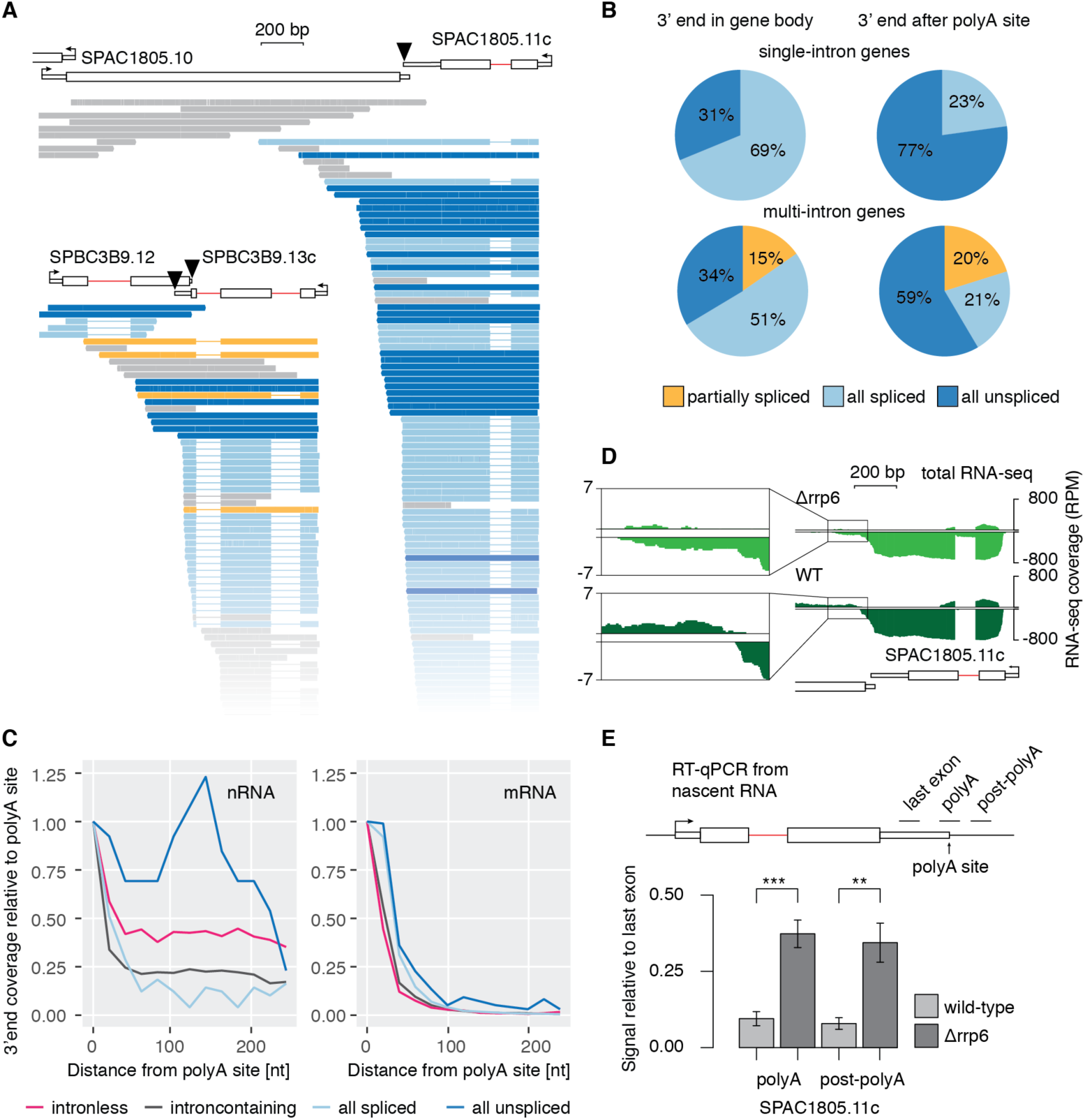
Coupling between co-transcriptional splicing, polyA site cleavage, and mRNA stability. (A) Examples of intron-containing genes with unspliced transcripts extending over the polyA site. Black triangles mark the polyA cleavage sites. Full representation of all sequenced transcripts in Fig. S8B. (B) Pie charts reflecting the fraction of spliced and unspliced transcripts in single or multi-intron genes with 3’ ends either within the gene body or downstream of the polyA site cleavage site. (C) 3’ end profiles downstream of annotated polyA cleavage sites for different transcript classes. Data are binned in 20 nt intervals and normalized to first bin (-20 nt – 0 nt from polyA cleavage site). (D) WT and Δ*rrp6* total RNA-seq read coverage over the same example gene as in A is shown (log-scale). Counts per nucleotide were normalized to library size. Data reanalyzed from (Zhou et al. 2015). The inset zooms into the region downstream of the annotated polyA site. (E) Nascent transcript levels with 3’ ends extending over the polyA site are increased in the exosome mutant Δ*rrp6*. RT-qPCR from Δ*rrp6* and WT strains confirmed higher levels of nascent RNA uncleaved at the polyA cleavage site, using qPCR primers to generate amplicons (black line above gene diagram) bridging (polyA) or downstream (post-polyA) of the polyA cleavage site (RT with random hexamers). SDs from 4 biological replicates are given. Asterisks indicate significance according to the Student’s t-test (p < 0.05 *, p < 0.01 **, p < 0.001 ***).

## Discussion

In this study, we sought to determine whether rapid co-transcriptional splicing in *S. pombe* enforces the removal of introns in the order of their transcription. In depth analysis of nascent RNA by short- and long-read sequencing revealed that distinct populations of fully spliced, fully unspliced and partially spliced nascent RNA are uniquely detectable by LRS. The preponderance of fully spliced nascent transcripts indicates that splicing proceeds rapidly and efficiently in the order of intron synthesis with few exceptions. Moreover, we show that fully unspliced nascent transcripts, which account for the majority of unspliced introns, fail to terminate properly and are degraded in the nucleus. This evidence extends previous co-transcriptional splicing studies from multiple species. Specifically, bulk determinations by short-read RNA-seq yield co-transcriptional splicing efficiencies on a per intron basis, ranging from 45-84% depending on the species and platform (Brugiolo et al. 2013; Herzel and Neugebauer 2015). It is generally assumed that remaining introns are post-transcriptionally spliced. Instead, our findings indicate that, at least in *S. pombe*, the majority of introns that are not spliced co-transcriptionally will be degraded. We demonstrate that degradation of unspliced RNA contributes significantly to shaping the transcriptome in response to cellular stimuli changing gene expression. An apparent cooperativity among neighboring introns may lead to “all or none” splicing of individual transcripts, suggesting a specific mechanism(s) for crosstalk among spliceosomes and the polyA site cleavage machinery during Pol II elongation.

Our analysis highlights intensive chromatin-associated splicing activity in *S. pombe*. Mass spectrometry of the isolated chromatin fraction identified numerous splicing factors that were not detected in *S. cerevisiae* chromatin (Carrillo Oesterreich et al. 2010). This likely reflects the 10x higher number of intron-containing genes in the *S. pombe* genome compared to *S. cerevisiae* and is consistent with co-transcriptional splicing and/or the retention of post-splicing intron lariat spliceosomes on chromatin (Chen et al. 2014). Analysis of splicing frequency on a per intron basis from nascent RNA-seq showed that most *S. pombe* introns are spliced co-transcriptionally (median 0.59), which is lower than in *S. cerevisiae* (median 0.74), *Drosophila* or human cells (Brugiolo et al. 2013). Notably, 5% of mRNAs contain introns; recent studies suggest that translation of retained introns contributes to protein diversity in *S. pombe* (Duncan and Mata 2014; Kuang et al. 2017). Intron length is often associated with reduced co-transcriptional splicing (Carrillo Oesterreich et al. 2010; Brugiolo et al. 2013). However, we (see Figure 2A and S2C) and others (Eser et al. 2016) find that intron length is one of the few parameters that does *not* correlate with nascent RNA splicing frequency, while 5’SS and BPS strength do. Low co-transcriptional splicing of first introns in *S. pombe* and other species suggests that the first intron can persist while downstream introns are spliced (Kessler et al. 1993; Khodor et al. 2011; Khodor et al. 2012; Tilgner et al. 2012).

Full-length nascent transcript sequences reveal the life history of individual (pre-)mRNAs that cannot be inferred from conventional short-read sequencing. Data on full-length transcripts can be obtained by RT-(q)PCR (Kessler et al. 1993; Pandya-Jones and Black 2009; Glauser et al. 2011; Bonde et al. 2014), synthetic long-read sequencing (Tilgner et al. 2015; Tilgner et al. 2018) or LRS on the Oxford Nanopore and Pacific Biosciences platforms (Byrne et al. 2017). The latter platform has been previously used to characterize full-length mRNA splicing products (Sharon et al. 2013; Kuang et al. 2017). Here, we used it to analyze splicing substrates: full-length nascent transcripts. The mapping of nascent RNA 3’ ends reports on Pol II position within genes. Promoter-proximal Pol II pausing was identified in *S. pombe* (Booth et al. 2016) and NET-Seq was recently used to analyze transcription elongation (Shetty et al. 2017). Although higher coverage will be required to investigate Pol II dynamics, LRS of nascent RNA represents an additional powerful approach for investigating RNA processing and transcription regulation.

Full length sequencing of individual nascent RNAs enabled us to determine the fate of unspliced transcripts. Most often, unspliced nascent transcripts fail to undergo polyA site cleavage, leading to read-through transcription and RNA degradation by the nuclear exosome. Therefore, our data complement prior reports showing that unspliced transcripts are degraded in an *rrp6*-dependent manner (Bousquet-Antonelli et al. 2000; Bitton et al. 2015; Zhou et al. 2015; Kilchert et al. 2016). Our work does not exclude the action of additional degradation pathways for unspliced transcripts, like the nuclear 5’-3’ exonuclease Dhp1/Rat1/Xrn2 and cytoplasmic nonsense mediated decay (Vargas et al. 2011; Davidson et al. 2012; Girard et al. 2012; Kervestin and Jacobson 2012; Brogna et al. 2016; Kilchert et al. 2016). Physical interactions have been detected between spliceosomal components and the nuclear decay machinery to provide a layer of surveillance based on splicing fidelity (Nag and Steitz 2012; Zhou et al. 2015; Herzel et al. 2017). Our findings suggest that these interactions take place co-transcriptionally, explaining why poor co-transcriptional splicing is associated with lowered mRNA levels independent of transcriptional changes upon caffeine treatment (this study) and possibly why splicing levels within mRNA correlate with expression levels (Wilhelm et al. 2008). Cooperativity could drive the formation of fully spliced mRNAs over partially spliced transcripts to evade decay in the nucleus or cytoplasm. Also, increased mRNA half-life was observed *in vitro* when splicing occurred in a transcription-coupled system (Hicks et al. 2006).Conversely, fully unspliced transcripts might be degraded more efficiently. Perhaps these transcripts arise during the process of shutting down gene expression. For example, read-through transcription and inhibition of splicing can occur during mitosis, cellular senescence, and/or under conditions of cellular stress (Leong et al. 2014; Shalgi et al. 2014; Vilborg et al. 2015; Muniz et al. 2017), though read-through transcription has not yet been linked to splicing inhibition in these cases. Taken together, the link between co-transcriptional splicing and mRNA half-life is an important determinant of gene expression.

Is cooperativity among splicing events widespread? We found that a single spliced or unspliced intron can mediate positive and negative effects on the splicing of other introns within the transcript. Our results *in vivo* are completely consistent with previous *in vitro* experiments showing that splicing of multi-intron transcripts is more efficient following an initial intron removal event (Crabb et al. 2010). In *S. cerevisiae*, constitutive splicing of the 2-intron *SUS1* gene has been found to occur cooperatively (Bonde et al. 2014). Co-association, the likelihood that distinct splicing events are correlated, has been detected for multiple alternative splicing events in human cells, *C. elegans* and *S. pombe* (Fededa et al. 2005; Glauser et al. 2011; Tilgner et al. 2015; Kuang et al. 2017; Tilgner et al. 2018). Importantly, a recent study concluded that splicing is regulated by adjacent processing events, based on deduced order of intron removal from human paired end RNA-seq (Kim et al. 2017). In the context of alternative splicing, intron removal can be delayed relative to an intron downstream of a cassette exon (Kim et al. 2017); interestingly, these delayed introns can already be committed to splicing (de la Mata et al. 2010). Thus, this under-appreciated phenomenon of splicing co-association has been observed in multiple species in total RNA. Here, it is extended to full-length nascent RNA, detecting intermediate states in RNA processing before and after splicing and enabling further mechanistic insights into how crosstalk is mediated.

Cooperativity among splicing events and with polyA cleavage likely occurs at the level of assembling spliceosomes. Median spliceosomal protein copy number per cell is almost identical to the number of intron-containing genes in *S. pombe* (Marguerat et al.2012). Hence, high local concentrations of splicing factors at sites of active transcription might promote splicing when the splicing machinery is at non-saturating levels or in the context of competition with RNA decay (Crabb et al. 2010; Munding et al. 2013). The cellular concentration of polyadenylation factors has also been linked to coupled alternative polyadenylation and splicing of the terminal intron (Movassat et al. 2016). Additional features contributing to cooperativity may include the local chromatin environment and/or changes in Pol II elongation and modifications (Gunderson and Johnson 2009; Gunderson et al. 2011; Patrick et al. 2015; Milligan et al. 2017; Neves et al. 2017; Nissen et al. 2017). An EM tomography study of the Balbiani ring 3 gene containing 38 introns suggested that only one spliceosome may associate with each nascent RNA at any given time (Wetterberg et al. 2001). While this remains to be tested in other systems, spliceosomal components can remain associated with multi-intron pre-mRNAs during splicing (Crabb et al. 2010; de la Mata et al. 2010). This would be consistent with the possibility that the presence of pre-spliceosomal components, assembling spliceosomes, or remaining spliceosomal components after splicing may enhance or antagonize cooperativity among introns (Huang et al. 2002; Crabb et al. 2010; Chen et al. 2014). In mammalian cells, U2AF interacts directly with the polyA cleavage machinery (Vagner et al. 2000; Millevoi et al. 2006), while U1 snRNP interactions with nascent RNA – particularly in first introns – suppress premature polyA cleavage, prevent polyadenylation, and enhance transcription elongation through the gene’s entirety (Gunderson et al. 1998; Berg et al. 2012; Park and Hannenhalli 2015; Oh et al. 2017; Chiu et al. 2018). Perhaps U1 snRNP lingering on unspliced introns can promote elongation and inhibit cleavage at correct polyA sites as well, producing the observed read-through transcription. The overrepresentation of unspliced first introns in all organisms tested leads us to speculate that splicing of the first intron represents a tipping point in a transcript-specific decision of whether to proceed with transcription, splicing, and 3’ end formation.

## Methods

***S. pombe* strains, genome version and annotation, growth conditions**

For strain information please refer to Table S1. The *S. pombe* genome version EF2 (http://goo.gl/PuaBni) and gene annotation ASM294v2.31 were used (Eser et al. 2016) (Table S5). *S. pombe* cell cultures were grown in complete liquid media (YES) at 30°C and 250 rpm. The ***Δ****rrp6* strain was grown under G418 selection. Cells were harvested by centrifugation in exponential growth at OD (595nm) ∼0.5. Cell pellets were washed with ice-cold PBS, quick-frozen in liquid nitrogen in 6 aliquots and kept at −80°C. For *prp2-1* splicing analysis YES cultures originating from single *prp2-1* and 972h cell colonies were grown at 26°C to an OD of 0.4, then shifted to 37°C for 2 hours before harvesting.

### Protein analysis

For western blotting, ∼3 μg of protein were loaded per lane. The following antibodies were used: Rpb1 (8WG16), Histone H3 (ab1791, Abcam), GAPDH (Novus Biologicals, NB300-221) and ribosomal protein L5 (Santa Cruz, sc-103865). Coomassie gel staining and mass spectrometry were performed as described (Shevchenko et al. 2006) at the mass spectrometry facility of MPI-CBG Dresden (Supplemental methods).

### RNA preparation

Nascent RNA was prepared as described (Carrillo Oesterreich et al. 2016) and used for nascent RNA sequencing and long-read sequencing (Supplemental methods). PolyA-RNA was obtained using oligo-dT coated magnetic beads binding to polyA+ RNA (Dynabeads mRNA DIRECT Micro Purification Kit, Life technologies). rRNA was removed from ∼5 μg chromatin-associated polyA-RNA using the Ribo-Zero Gold rRNA Removal Kit (Yeast) from Epicentre/Illumina.

### Qualitative and quantitative analysis of nucleic acids

RNA and DNA samples were analyzed by agarose (1-1.5%) or TBE-Urea polyacrylamide (10 or 15%, Invitrogen) gel electrophoresis. DNA and RNA concentrations were determined by UV/Vis spectroscopy with the Nanodrop2000 (ThermoScientific) or fluorometric measurements with the Qubit dsDNA BR Assay or the RNA BR Assay (Life technologies). Reverse transcription of RNA was done using SuperScript III reverse transcriptase (Invitrogen) with random hexamers, oligo(dT) or gene-specific primers. For details on RT, qPCR and PCR protocols see the Supplemental methods.

### Short-read RNA sequencing

For RNA-seq of different cellular fractions RNA samples were submitted to the Yale Center for Genome Analysis (YCGA). PolyA+ RNA depleted, rRNA depleted nascent RNA and cytoplasmic polyA+ RNA were analyzed by single-end RNA-seq with 76bp read length. Random hexamer primed Illumina libraries were prepared with standard protocols.

### Mapping short-read RNA-seq data to the *S. pombe* genome

Data were mapped to the genome with Tophat2 (version 2.0.12) (Kim et al. 2013) using the following settings: tophat2 -p 5 -i 30 -I 900 –g 1 -N 2 -G <Spombe_EF2> –segment-length 25 –library-type fr-firststrand –minanchor-length 8 –splice-mismatches 0 –min-coverage-intron 30 –max-coverageintron 900 –min-segment-intron 30 –max-segment-intron 900. RNA-seq coverage was adjusted for library size, excluding reads mapped to rDNA regions, weighted using the R-package edgeR (calcNormFactors without additional parameters, further details in Supplemental methods). To determine gene expression values between replicates and different samples cufflinks version 2.2.1 and cuffdiff were used with the following settings: cufflinks -p 20 -G <Spombe_EF2> -b <Spombe_EF2.fasta>; cuffdiff –frag-bias-correct <Spombe_EF2.fasta> –num-threads 20 – library-type fr-firststrand –library-normmethod geometric <Spombe_EF2.gtf>.

### Intron splicing quantification

For intron splicing calculation junction reads originating from spliced (split, cigar contains “N” and unspliced (unsplit, cigar contains “M“, but no “N”) transcripts were extracted from all mapped reads. Overlaps with 8 nucleotide windows around annotated 5‘ SS and 3‘ SS junctions were searched for split and unsplit reads separately with bedtools intersect. An overlap of at least 4 nucleotides on each side of the junction was required. The sum of all split reads per 5’SS (identical with 3’SS split read count) was divided by the sum of the 5’SS split read count and the average of unsplit reads over 5’ and 3’SS per intron. This resulted in a splicing score (splicing per intron, SPI) ranging from 0 to 1 with 1 being 100% spliced (Herzel and Neugebauer 2015). A cutoff of at least 10 reads per junction to report an SPI was applied.

### 3’ end ligation and long-read sequencing library preparation

3’ end ligation and long-read sequencing library preparation of polyA+ and rRNA-depleted, chromatin-associated RNA was done as described previously (Carrillo Oesterreich et al. 2016). Sample-specific barcodes were included in the PCR oligonucleotides (Table S2) for the 3 *prp2-1* cDNA libraries and the corresponding 972h-cDNA libraries. Double-stranded cDNA (>1 μg) was submitted to the Yale Center for Genomic Analysis (YCGA) for Pacific Biosciences library preparation and sequencing with standard protocols (SMRTbell Template Prep Kit 1.0). Sequencing was done with either diffusion- or magbead-loading (further details in Supplemental methods).

### Long-read sequencing data processing and mapping

3’ end linker sequences, Clontech adaptor sequences (SMARTer cDNA synthesis kit, Clontech) and the 5 nt random 3’ barcode were removed with cutadapt (Martin 2011) and the FASTX toolkit (http://hannonlab.cshl.edu/fastx_toolkit/index.html). Processed reads were mapped to the respective genome using gmap (Wu and Watanabe 2005). To remove potential mRNA contaminants reads ending within +/-100 nt of an annotated polyA site and short polyA tails (> 4 nt) were removed from the dataset.

### Classification of multi-intron transcripts

Reads overlapping completely with multiple introns were considered multi-intron transcripts. If the number of exons in a read was identical to the number of overlapping introns +1, a read was considered to be “all spliced”. If the block count was 1 irrespective of the number of overlapping introns, a read was considered to be “all unspliced”. Reads, which had a block count greater 1, but fewer block counts than the number of overlapping introns +1, were classified as “partially spliced” (further details in Supplemental methods).

### Prediction of transcript splicing status from nRNA-seq

To determine the expectation of the abundance of partially spliced transcripts, in the case that individual intron splicing is independent of each other, we assume that the SPI, calculated from short read RNA-seq, reflects the probability of splicing of a particular intron in the chromatin fraction at steady state. Further, we assume that the gene expression values from nRNA-seq reflect an estimate of relative transcript number per gene in the chromatin fraction at steady state. For each intron ‘spliced’ or ‘unspliced’ sampling with replacement was performed according to its SPI and the average transcript number per gene. Simulated data were combined into a matrix with columns representing each intron and rows representing each transcript. If a row contained both ‘spliced’ and ‘unspliced’ column entries, it was classified as ‘partially spliced’. If a row contained only ‘spliced’ or only ‘unspliced’ column entries, it was classified as ‘all spliced’ or ‘all unspliced’, respectively. In the gene-specific analysis a numerical solution using SPIs as splicing probability was calculated (Supplemental methods). From the gene-specific observed and predicted fraction of “all or none” splicing we calculated a co-association score, which we define as the log2-ratio between observed and predicted fraction of “all or none” splicing.

## Data access

The accession number for the data reported in this paper is GEO: GSE104681. Multiplexed WT and *prp2-1* CCS reads can be downloaded from Harvard Dataverse doi:10.7910/DVN/PW1KEG (Herzel 2017).

## Acknowledgements

We thank Gene-Wei Li and members of the Neugebauer and Li labs, Hanspeter Herzel, Michael Weber, Fernando Carrillo Oesterreich and Charles Query for discussions and comments on the manuscript. We are grateful to Iva Tolic, Tamas Fischer and Charles Query for gifts of strains, Jeremy Schofield for help with preliminary experiments, and Guilin Wang and the Yale Center for Genome Analysis for technical assistance and advice. The work was supported by funding by NIH R01GM112766 from the NIGMS. Its contents are solely the responsibility of the authors and do not necessarily represent the official views of the NIH. LH is a postdoctoral fellow of the Helen Hay Whitney foundation.

## Author contributions

LH and KMN conceived the study and designed the experiments. Semi-quantitative RT-PCR experiments represented in Figure S2B were carried out by KS. All other experiments and data analyses were performed by LH. LH and KMN wrote the manuscript. All authors read and approved the manuscript.

## Disclosure declaration

The authors declare no conflict of interest.

